# Investigating the Mitochondria-Related Mechanisms of Ligustri Lucidi Fructus in Treating Diabetic Nephropathy: A Combined Approach of Network Pharmacology, Transcriptomics, and Experimental Validation

**DOI:** 10.64898/2026.01.08.698456

**Authors:** Qinqing Li, Jie Yang, Yanli Xin, Lixia Wen, Ruqiao Luan, Xuelan Zhang

## Abstract

Mitochondrial dysfunction and excessive oxidative stress within mitochondria are key pathological factors driving renal tubular injury in diabetic nephropathy (DN). While Ligustri Lucidi Fructus (LLF) is traditionally used in the treatment of DN, its precise mechanisms, particularly those involving mitochondria, remain poorly understood. This study identifies differentially expressed genes (DEGs) through differential expression analysis of the GSE142025 dataset, which includes data related to DN. Feature genes were selected from yielding overlapping genes by cross-referencing four distinct machine learning models. Genes with significant differential expression and consistent trends across both datasets were subjected to receiver operating characteristic curve analysis, and those with AUC > 0.7 in both datasets were defined as biomarkers. To investigate potential mechanisms of action, function enrichment, immune infiltration, network construction, and molecular docking analyses were performed. Finally, a DN mice model was constructed, further confirming the therapeutic effect of LLF on DN through assessments of blood glucose, urinary microalbumin levels, and histopathological examination. Meanwhile, the expression of these biomarkers was validated using RT-qPCR. The results indicated that four biomarkers were associated with mitochondria and related to the treatment of DN by LLF. These biomarkers were enriched in pathways related to ribosome function, valine, leucine, and isoleucine degradation, cytokine-cytokine receptor interactions, and peroxisomes. All biomarkers were negatively correlated with CD8^+^ T cells and activated mast cells while showing positive correlations with activated NK cells and naive B cells. Additionally, the binding energies of taxifolin, beta-sitosterol, and eriodictyol with the biomarkers were all below -5 kcal/mol. Animal experiments have confirmed that LLF significantly upregulates the expression of CAT and MAOA genes in kidney tissue. Identifying mitochondrial biomarkers for LLF treatment of DN offers novel insights into therapeutic strategies for the disease.

## Introduction

Diabetic nephropathy (DN) is the most prevalent complication of diabetes mellitus and the leading cause of end-stage renal disease globally[1]. It is characterized by the accumulation of extracellular matrix in both the glomerular and tubular compartments, along with thickening and sclerosis of intrarenal blood vessels [2]. DN is commonly associated with proteinuria and hypertension[3]. Its development is closely linked to vascular endothelial cell damage, an exacerbated inflammatory response, and heightened oxidative stress resulting from prolonged hyperglycemia [4]. The incidence of DN is rising worldwide, particularly among middle-aged and elderly diabetic individuals. As the disease progresses, it can lead to end-stage renal failure and even cardiovascular complications, significantly impairing patients’ quality of life and prognosis [5]. Despite advances in diagnostic and therapeutic approaches for DN, early and accurate biomarkers for diagnosis remain elusive, and effective treatment strategies to reverse the pathological processes are still lacking. Hence, there is an urgent need for the development of novel, targeted anti-DN drugs.

Mitochondria are critical in cellular bioenergetics, metabolic precursor synthesis, calcium homeostasis, reactive oxygen species (ROS) production, immune signaling, and apoptosis, all of which are essential for maintaining cellular and organismal stability [6]. As the cell’s powerhouses, mitochondria play a pivotal role in fundamental processes such as glycolysis, the tricarboxylic acid cycle, and oxidative phosphorylation [7]. Obesity disrupts the Krebs cycle and mitochondrial respiratory chain, leading to mitochondrial dysfunction and increased ROS production. Elevated ROS levels in the mitochondrial respiratory chain can induce oxidative stress, which exacerbates the inflammatory response associated with obesity and promotes apoptosis[8]. Recent research has highlighted the significant role of mitochondrial dysfunction in the pathogenesis and progression of DN, including disturbances in energy metabolism, excessive ROS generation, and heightened apoptosis signaling [9]. Chronic mitochondrial dysfunction accelerates kidney disease progression [10]. Thus, improving mitochondrial function could represent a crucial protective strategy against DN.

*Ligustri Lucidi Fructus* (LLF), a dried ripe fruit from the Luteaceae family, is renowned for its liver and kidney-nourishing properties, as well as its benefits in darkening hair and improving vision [11]. A naturally occurring heteropolysaccharide extracted from LLF has been identified, revealing its potential to protect the kidney from fibrosis[12]. In recent years, increasing attention has been given to the use of LLF in treating DN, demonstrating notable renal protective effects[12–14]. Moreover, the intricate relationship between LLF and mitochondria has been well-documented and validated through extensive research. A notable study has elucidated that LLF exerts its beneficial effects by modulating mitochondrial function via the activation of the AMPK signaling pathway[15]. This mechanism effectively fortifies mitochondria against damage caused by oxidative stress. Such insights further highlight the crucial role of LLF in sustaining cellular energy metabolism and enhancing the resilience of cells to oxidative challenges. However, the precise therapeutic mechanism, particularly the role of mitochondrial function recovery, remains inadequately understood.

This study aimed to elucidate the biological mechanisms of the therapeutic effect of LLF on mitochondrial function in DN. By integrating transcriptomic data and active ingredient information from public databases, bioinformatics tools were employed to identify biomarkers linked to the renal protective effects of LLF. Further analyses, including immune infiltration, clinical feature association, m^6^A RNA modification, functional enrichment, regulatory network construction, and molecular docking, suggested that these biomarkers play a pivotal role in regulating mitochondrial function during DN treatment. *In vivo* validation further confirmed their significance. This comprehensive analysis not only deepens the understanding of mechanisms of LLF in treating DN, but also provides a solid foundation for developing novel therapeutic targets based on mitochondrial dysfunction.

## Materials and methods

### Data collection

The gene expression matrix and corresponding clinical data for the GSE142025 and GSE96804 datasets related to DN were retrieved from the Gene Expression Omnibus (GEO) database (https://www.ncbi.nlm.nih.gov/geo/) [16]. GSE142025, used as the training set, included kidney tissue samples from 27 patients with DN and 9 controls, sequenced using the GPL20301 platform. GSE96804, serving as the validation set, comprised sequencing data from 41 patients with DN and 20 controls, processed with the GPL17586 platform. Both datasets focus on kidney tissue, with the GSE96804 dataset specifically examining the glomerulus, the primary filtration unit of the kidney(Fig. 1).

**Fig. 1.**
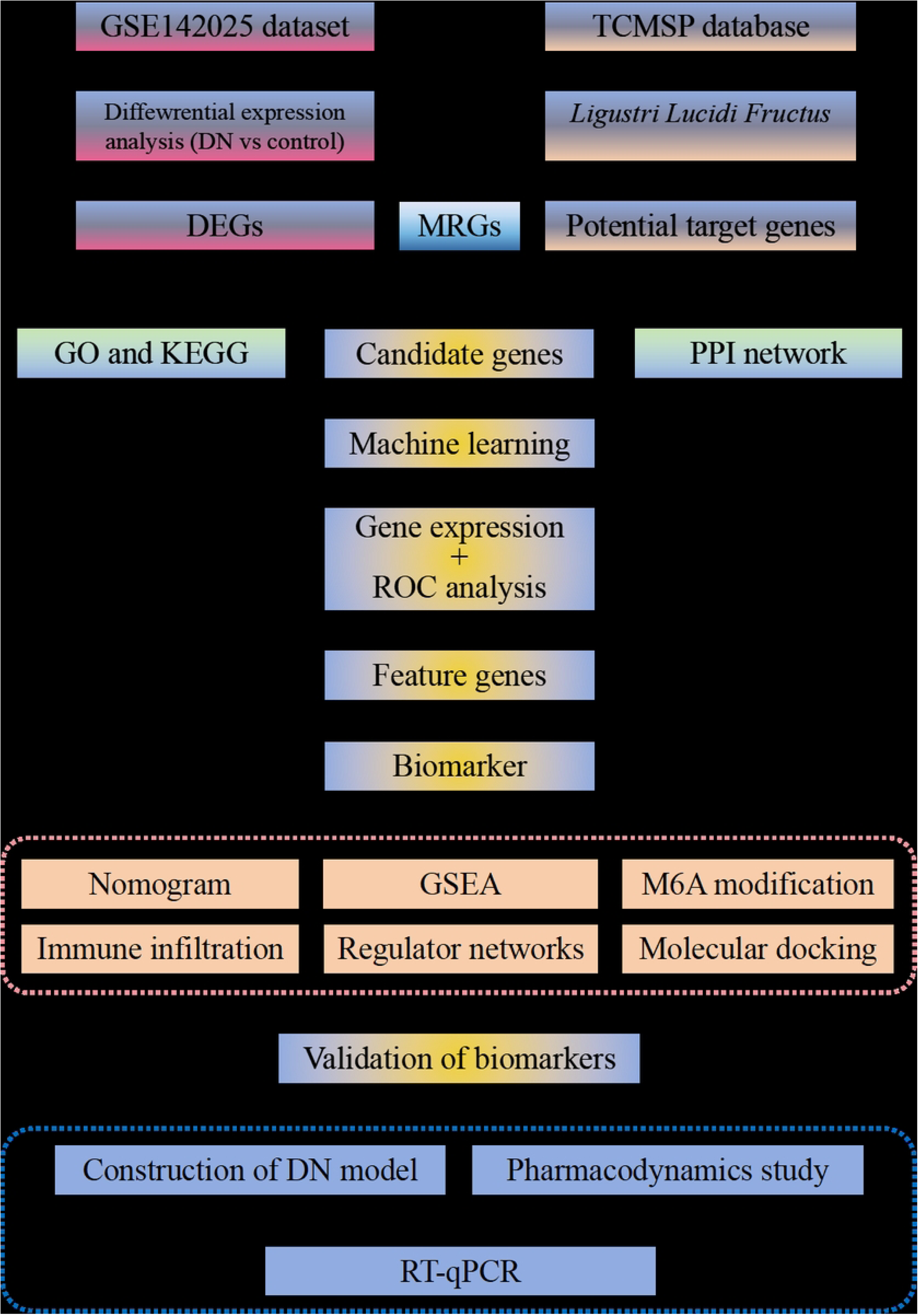
Flowchart of the Study Design

A total of 1136 mitochondria-related genes (MRGs) were extracted from the MitoCarta3.0 database (https://www.broadinstitute.org/mitocarta). The active ingredients of LLF were predicted using the Traditional Chinese Medicine Systems Pharmacology (TCMSP) database (http://sm.nwsuaf.edu.cn-/lsp/tcmsp.php), based on an oral bioavailability (OB) threshold of ≥30% and a drug-likeness (DL) threshold of ≥ 0.18 [17]. Subsequently, potential target genes for the active constituents were predicted using the Swiss Target Prediction database (http://www.swisstargetpred-iction.ch/).

### Differential expression analysis

Differential expression analysis of GSE142025 (DN vs. control) was performed using the limma package (v 3.54.1), with significance criteria set at *P*.adj < 0.05 and |log2FoldChange| > 0.5 [18]. Volcano plots and heat maps were visualized using the ggplot2 (v 3.3.6) and ComplexHeatmap (v 2.14.0) packages, respectively [19–20]. DEGs, MRGs, and potential target genes of the active ingredients were intersected, and the overlapping genes were defined as candidate genes. The network linking the active ingredients and candidate genes was constructed using Cytoscape software (v 3.9.0) [21].

### Functional enrichment analysis and protein-protein interaction (PPI) network construction

Gene Ontology (GO) and Kyoto Encyclopedia of Genes and Genomes (KEGG) enrichment analyses for candidate genes were performed using the clusterProfiler package (v 4.6.2) to explore their biological functions and associated signaling pathways (*P*.adjust < 0.05). The candidate genes were then input into the STRING database (https://cn.string-db.org/) to retrieve PPI relationships (confidence level ≥ 0.4), followed by the construction of a PPI network using Cytoscape (v 3.9.0) [22].

### Machine learning

Four machine learning models—random forest (RF), k-Nearest Neighbor (KNN), Partial Least Squares (PLS), and support vector machine (SVM)—were developed using the caret package (v 6.0-93) with data from GSE142025 [https://github.com-/topepo/caret/]. The DALEX package(v2.4.3) [https://github.com/-ModelOriented/DALEX] was employed to analyze the models and generate residual distribution plots. The importance of candidate genes was assessed using root mean square error (RMSE). A smaller RMSE value indicates higher predictive accuracy of the model[23–24]. In our study, based on the data, we selected a threshold RMSE value of 0.281. Genes exhibiting an RMSE value below this threshold across all four models were selected as feature genes for further analysis.

### Identification of biomarkers

In GSE142025 and GSE96804, the Wilcoxon test was applied to compare the expression of feature genes between DN and control samples. Genes exhibiting significant expression differences (*P* < 0.05) and consistent expression trends across both datasets were selected for receiver operating characteristic (ROC) curve analysis. The pROC package (v1.18.0) was used to generate ROC curves and calculate the area under the curve (AUC), with genes showing AUC > 0.7 in both datasets classified as biomarkers [25].

### Gene set enrichment analysis (GSEA)

To further investigate the biological functions and signaling pathways associated with the biomarkers, GSEA was performed on the GSE142025 dataset. First, Spearman correlation analysis of biomarkers with all other genes was performed using the psych package (v2.2.9)[26]. Correlation coefficients were calculated and ranked (from high to low). The reference gene set used was c2.cp.kegg.v2023.1.Hs.symbols.gmt from the Molecular Signatures Database (MSigDB, https://www.gsea-msigdb.org/gsea/msigdb/). GSEA was then conducted evaluate the enrichment of ranked genes in the background gene set using the clusterProfiler package (v4.6.2). Multiple testing correction was applied *via* the FDR method, and adjusted *P-*values (denoted as *P*.adjust) were considered significant if < 0.05.

### M6A modification analysis

To investigate RNA methylation modifications of biomarkers, the SRAMP database (http://www.cuilab.cn/sramp/) was utilized to predict m^6^A modification sites on biomarkers, focusing on high-confidence positions within their secondary structures. The ENCORI database (https://starbase.sysu.edu.cn/) was then employed to identify m6A-modified proteins interacting with biomarkers, with the parameter |HepG2 (shRNA)| > 1 set for screening key proteins. The RPISeq database (http://pridb.gdcb.iasta-te.edu/RPISeq/) was subsequently used to predict the likelihood of interactions between key proteins and biomarkers. RNA sequences for both were uploaded in plain text format to generate RF and SVM classifier prediction scores. An interaction was considered significant when the score exceeded 0.5 [27].

### Immune filtration analysis

The CIBERSORT algorithm was applied to estimate the proportions of 22 immune cell types in both control and DN samples from GSE142025, with visualization achieved through heatmap generation using the ggplot2 package (v3.3.6) [28]. The Wilcoxon test was used to compare immune cell proportions between the two groups, and immune cells with *P* < 0.05 were designated as differential. Spearman correlation analysis between differential immune cells and biomarkers was performed using the psych package.

### Construction of networks and Molecular docking

MicroRNAs (miRNAs) interacting with biomarkers were predicted using the miRNet database (https://www.mirnet.ca). Subsequently, long noncoding RNAs (lncRNAs) targeting the identified miRNAs were predicted through the TarBase (http://www.diana.pcbi.upenn.edu/tarbase) and starbase (http://starbase.sysu.edu.cn/) databases. lncRNAs common to both databases were selected for network construction. A lncRNA-miRNA-mRNA regulatory network was then built using Cytoscape software. Core active ingredients targeting biomarkers were chosen to construct an active ingredient-biomarker network. Additionally, active ingredients, biomarkers, and pathways identified in GSEA were incorporated into Cytoscape to create an active ingredient-biomarker-pathway network.

Molecular docking analysis was performed to assess the binding affinity between core active ingredients and biomarkers. The 3D structures of biomarker proteins were obtained from the Research Collaboratory for Structural Bioinformatics Protein Data Bank (RCSB PDB, http://www.rscb.org/pdb) in PDB file format. The 2D structures of core active ingredients were retrieved in SDF format from the PubChem database (http://pubchem.ncbi.nlm.-nih.gov). Molecular docking was conducted using the CB-Dock platform (http://clab.labshare.cn/cb-dock/php/blinddock.php). A binding energy of less than -5 kcal/mol indicated strong binding affinity [29].

### Animal experiments

Twelve SPF-grade male db/db mice, aged 8-9 weeks, and six age-matched db/m mice were purchased from Changzhou Kavins Experimental Animal Co., Ltd. The mice were housed in the SPF animal facility of Shanxi University of Traditional Chinese Medicine and underwent a 7 day acclimatization in the laboratory under a 12/12 hour light cycle with ad libitum access to food and water. The research was approved by the Ethics Committee of Shanxi University of Traditional Chinese Medicine (2022DW167).

Following the acclimatization period, the establishment of a diabetic nephropathy (DN) model in db/db mice was confirmed by a tail vein blood glucose level ≥16 mmol/L, and the presence of microalbuminuria as indicated by a positive urine microalbumin test strip. Upon successful DN model establishment, the db/db mice were randomly divided into two groups (n=6 per group): the DN model group (DN) and the LLF treatment group (Treatment). Moreover, db/m mice (n=6) were used as the control group (Control). The Control and DN groups were given distilled water, whereas the treatment group received 3.5 g/kg LLF, for 8 weeks. After 8 weeks of administration, all mice were euthanized for serum, urine, and kidney tissue collection for subsequent testing.

### Blood and urine indicators

The serum levels of glucose were analyzed by a fully Automatic Blood Biochemistry Analyzer (Biobase, China, BK-280). The urinary microalbumin concentration was detected according to the instructions of the kit.

### Pathological observation of mouse kidney tissues

The kidney tissues were fixed in a 4% paraformaldehyde solution, followed by washing, dehydration, paraffin embedding, and sectioning. The paraffin-embedded kidney tissue sections were stained with hematoxylin and eosin (HE). Pathological changes in the kidney tissues were observed under an optical microscope.

### Reverse transcription quantitative polymerase chain reaction (RT-qPCR)

To verify biomarker expression, kidney tissue samples were collected from mice for RT-qPCR analysis. Total RNA was extracted from the samples using TRIzol reagent (Vazyme, China, R401-01) according to the manufacturer’s instructions. Notably, the concentration of the extracted RNA was measured using the NanoPhotometer N50, and the quality of the extracted RNA was assessed (Table 1). cDNA was synthesized from the RNA using the SureScript First-Strand cDNA Synthesis Kit (Servicebio, Wuhan, China). RT-qPCR analysis was performed using 2x Universal Blue SYBR Green qPCR Master Mix (Servicebio). Primer sequences for PCR are listed in Table 1, with GAPDH serving as the internal reference gene. Biomarker expression levels were calculated using the 2^−ΔΔCt^ method [14].

**Table 1.**
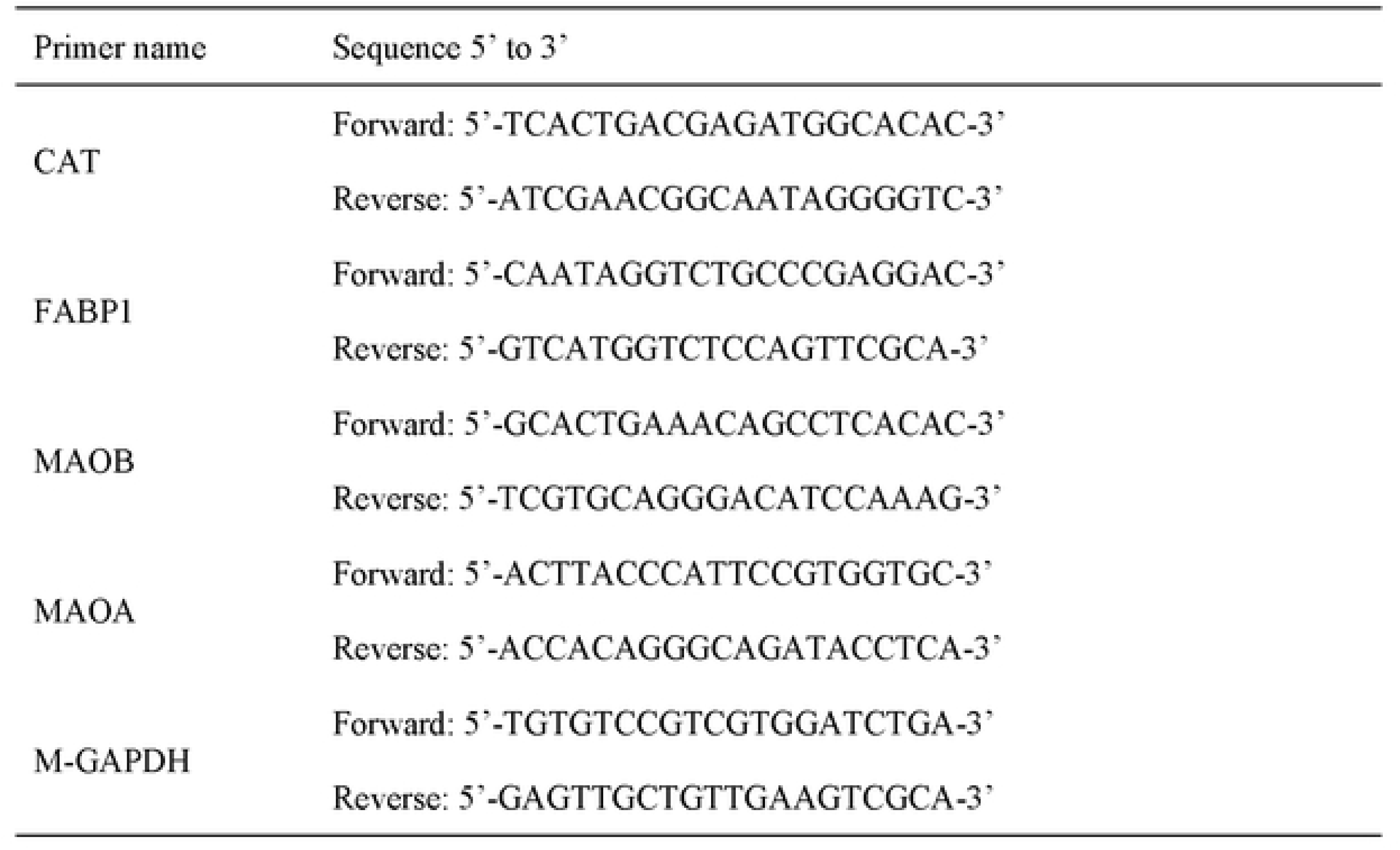
Primer sequences.

### Statistical analysis

Data processing and statistical analysis were performed using R software (version 4.2.2), with all methods selected and applied in accordance with statistical requirements. Differences between groups were assessed using the Wilcoxon test (*P* < 0.05). In our study, *** represents *P* <0.001, ** represents *P* < 0.01, * represents *P* < 0.05, and ns represents *P* > 0.05.

## Results

### Screening of candidate genes for LLF treating DN

In the GSE142025 dataset, 3,810 DEGs were identified between the DN and control groups, including 1,904 upregulated and 1,906 downregulated DEGs (Fig. 2A, B). Thirteen active ingredients from LLF were predicted using the TCMSP database, namely beta-sitosterol, kaempferol, taxifolin, Lucidumoside D, Lucidumoside D_qt, (20S)-24-ene-3,20-diol-3-acetate, eriodictyol, syringaresinol diglucoside_qt, Lucidusculine, Olitoriside, Olitoriside_qt, luteolin, and quercetin (Table 2). Four active ingredients—Lucidumoside D_qt, (20S)-24-ene-3,20-diol-3-acetate, syringaresinol diglucoside_qt, and Olitoriside_qt—did not predict any potential target genes, while the remaining nine ingredients predicted 517 potential target genes. By overlapping the 3,810 DEGs, 1,136 MRGs, and 517 potential target genes, nine candidate genes were identified: *GPX1*, *BAX*, *CASP8*, *MAOA*, *MAOB*, *CAT*, *AKR1B10*, *ALDH2*, and *FABP1* (Fig. 2C). An active ingredient-candidate gene network was subsequently constructed (Fig. 2D). These nine candidate genes were enriched in 341 GO terms, including response to toxic substances, organic hydroxy compound catabolic process, and cellular detoxification (Fig. 2E). Additionally, they were associated with 52 KEGG pathways, such as tryptophan metabolism, neurodegenerative pathways, and histidine metabolism (Fig. 2F).

**Fig. 2.**
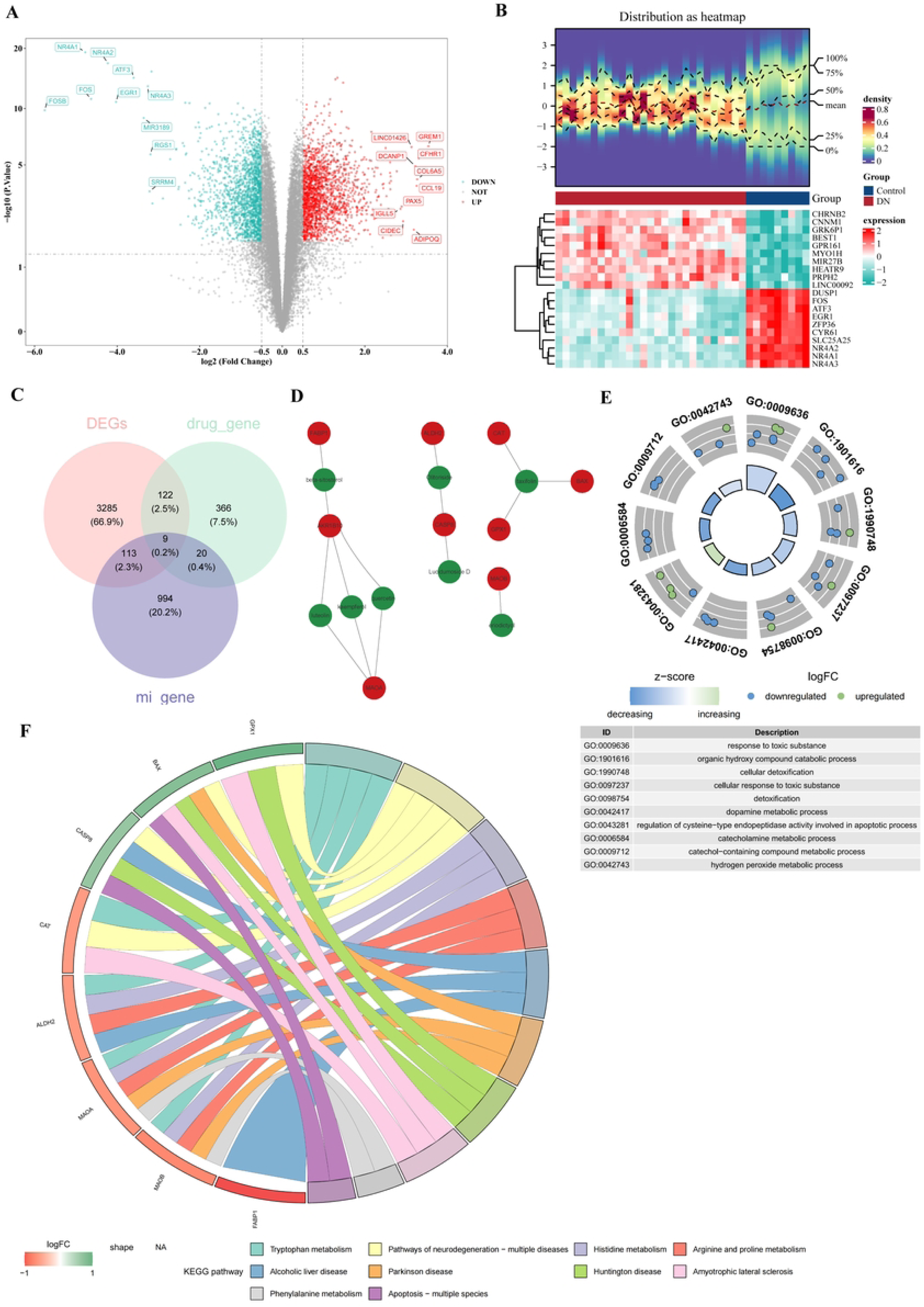
Screening of candidate genes. A Volcano plot showing differential gene expression between the DN and control groups. B Heatmap of top 10 up-regulated and down-regulated differential genes (ranked by |log2FC| from high to low) expression patterns between the DN and control groups. C Venn diagram illustrating the overlapping genes among DEGs, MRGs, and potential drug target genes. D Network of active ingredients and candidate genes. E Enrichment of candidate genes in GO terms. The left inner circle displays a bar chart, where the height of each bar indicates the significance of the corresponding term, with taller bars representing higher significance. The color of the bars reflects the z-score: darker colors signify higher z-scores, suggesting a greater likelihood of upregulation or downregulation of the biological function. The outer circle presents a scatter plot that shows the expression levels of each gene within each term, with green and blue dots representing upregulated and downregulated genes, respectively. The right side provides detailed descriptions of the GO enrichment terms. F KEGG pathway enrichment analysis for candidate genes.

**Table 2.** 13 Fructus Ligustris active ingredients.

| MOL ID | Molecule name | OB(%) | DL | Target number |
| --- | --- | --- | --- | --- |
| MOL000358 | beta-sitosterol | 36.91 | 0.75 | 100 |
| MOL000422 | kaempferol | 41.88 | 0.24 | 103 |
| MOL004576 | taxifolin | 57.84 | 0.27 | 92 |
| MOL005146 | Lucidumoside D | 48.87 | 0.71 | 104 |
| MOL005147 | Lucidumoside D_qt | 54.41 | 0.47 | 0 |
| MOL005169 | (20S)-24-ene-3,20-diol-3-acetate | 40.23 | 0.82 | 0 |
| MOL005190 | eriodictyol | 71.79 | 0.24 | 101 |
| MOL005195 | syringaresinol diglucoside_qt | 83.12 | 0.8 | 0 |
| MOL005209 | Lucidusculine | 30.11 | 0.75 | 105 |
| MOL005211 | Olitoriside | 65.45 | 0.23 | 100 |
| MOL005212 | Olitoriside_qt | 103.23 | 0.78 | 0 |
| MOL000006 | luteolin | 36.16 | 0.25 | 102 |
| MOL000098 | quercetin | 46.43 | 0.28 | 103 |

### Screening of biomarkers for DN treatment related to mitochondrial function in LLF

The PPI network revealed seven nodes and eight edges, with *MAOA*, *ALDH2*, *MAOB*, and *AKR1B10* interacting with each other (Fig. 3A). Genes with RMSE values less than 0.281 across four machine learning models were identified as feature genes: *CAT*, *MAOB*, *MAOA*, *BAX*, and *FABP1* (Fig. 3B-E). Expression analysis showed that *CAT*, *FABP1*, *MAOB*, and *MAOA* were significantly different between the DN and control groups and were consistent in both the GSE142025 and GSE96804 datasets (Fig. 3F, G). Furthermore, their AUC values in ROC curve analysis exceeded 0.7 in both datasets, indicating that these genes could effectively differentiate DN samples from control samples and serve as biomarkers for DN treatment related to mitochondrial function in LLF (Fig. 4A-H).

**Fig. 3.**
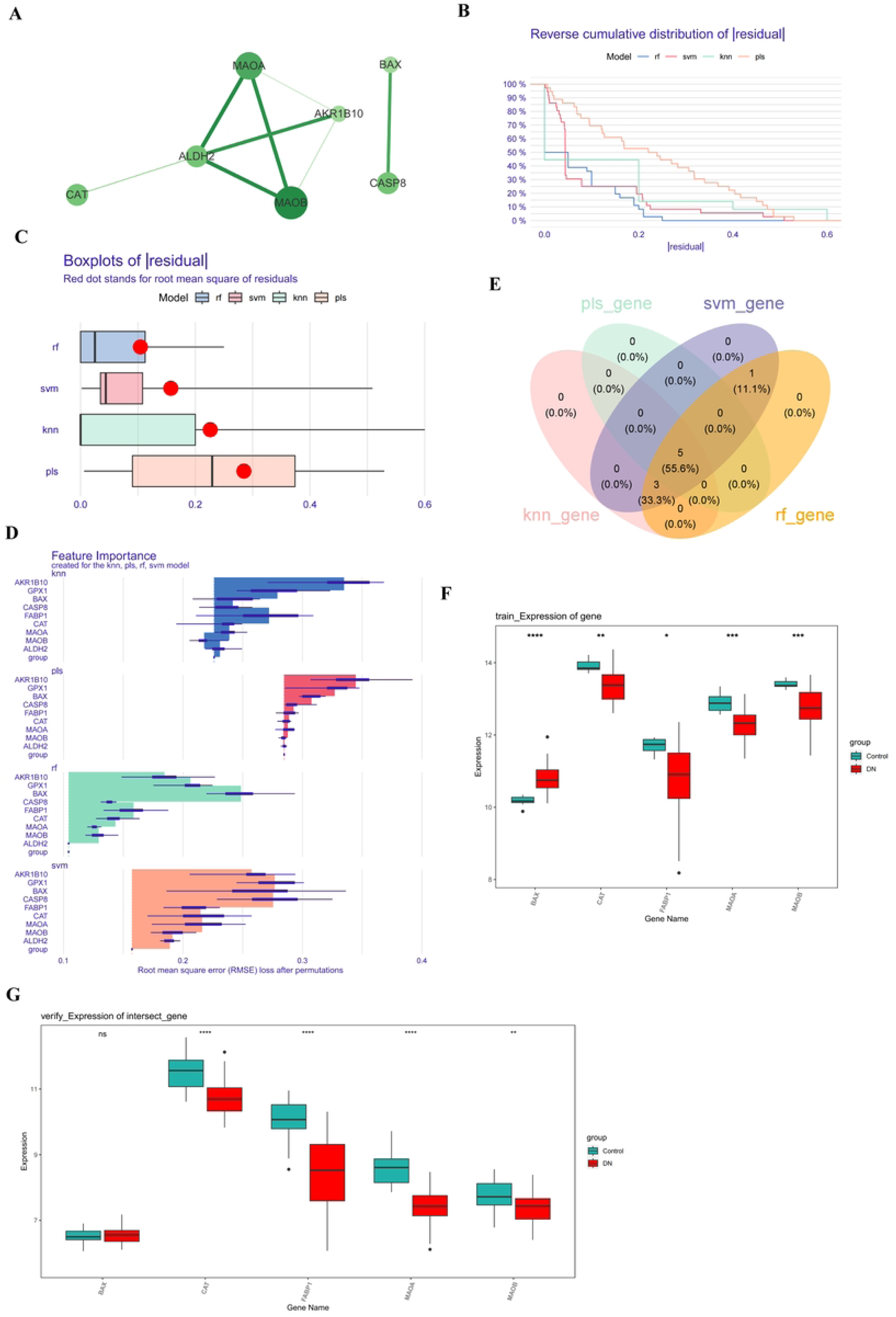
Four biomarkers. A PPI network of candidate genes. B Reverse cumulative distribution of residuals for four machine learning models: Random Forest (RF), k-Nearest Neighbor (KNN), Partial Least Squares (PLS), and Support Vector Machine (SVM). C Boxplots showing the distribution of residuals for the four models. The red dot indicates the root mean square of residuals. D Importance scores of candidate genes across the four machine learning models, assessed by comparing the RMSE (Root Mean Square Error) of different models. E Intersection of trait genes with RMSE < 0.281 across the four models, highlighting the common genes identified by all models. F-G Boxplots showing the expression levels of the four characteristic genes in GSE142025 and GSE96804 datasets. The x-axis represents the four genes, and the y-axis represents their expression levels. Green indicates the control group, and red represents the DN group.

**Fig. 4.**
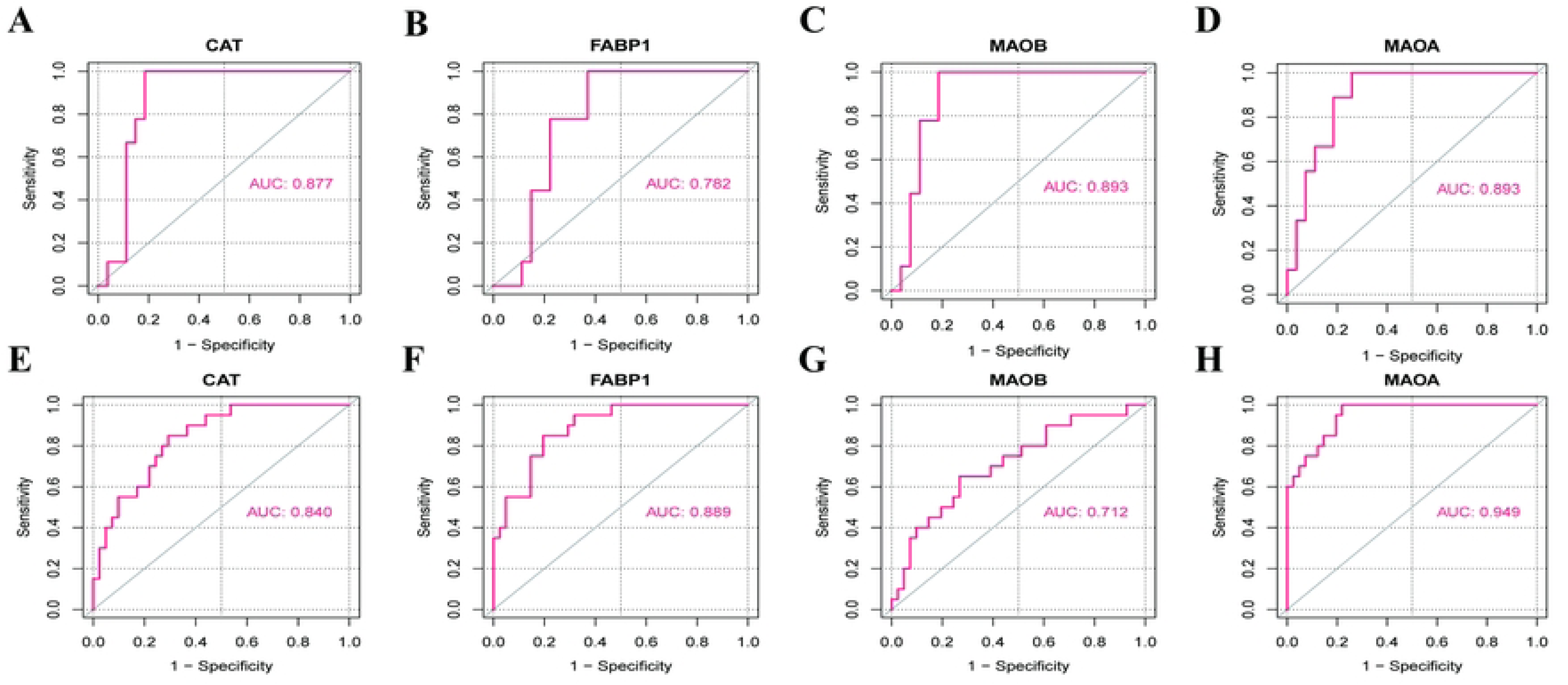
ROC curves for candidate genes. A-D Training set. E-H Validation set. The AUC value indicates the area under the curve.

### Biomarkers were significantly enriched in inflammatory and immune-related pathways

GSEA identified four biomarkers prominently enriched in the chemokine signaling pathway and cytokine-cytokine receptor interactions (Fig. 5A-D). Among these, the peroxidase signaling pathway showed a significant association with *CAT*, *MAOA*, and *MAOB*.

**Fig. 5.**
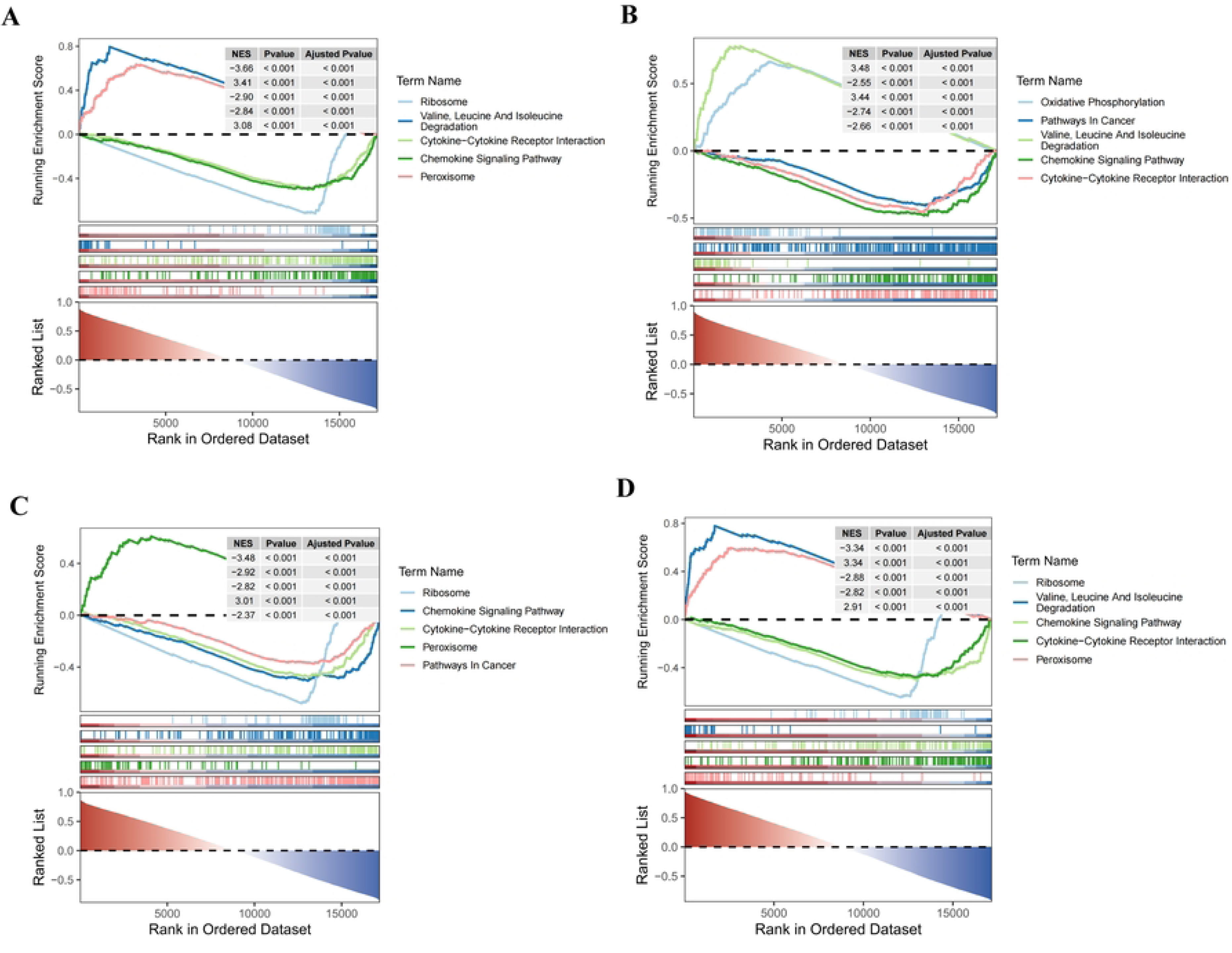
GSEA for biomarkers. A GSEA enrichment for CAT. B GSEA enrichment for FABP1. C GSEA enrichment for MAOA. D GSEA enrichment for MAOB.

### Biomarkers were correlated with immune cells

Notable differences in the expression of nine immune cell types-natrive B cells, M0 Macrophage, M1 Macrophage, M2 Macrophage, activated Mast cell, activated NK cell, resting memory CD4^+^ T cell, naive CD4^+^ T cell, and CD8^+^ T cell—were observed between the DN and control samples (*P* < 0.05) (Fig. 6A-B). A significant positive correlation (cor = 0.6) was found between naive B cells and activated NK cells, while a significant negative correlation (cor = -0.69) was detected between naive B cells and activated Mast cells (Fig. 6C). All biomarkers exhibited strong negative correlations with CD8^+^ T cells and activated Mast cells and positive correlations with activated NK cells and naive B cells (Fig. 6D).

**Fig. 6.**
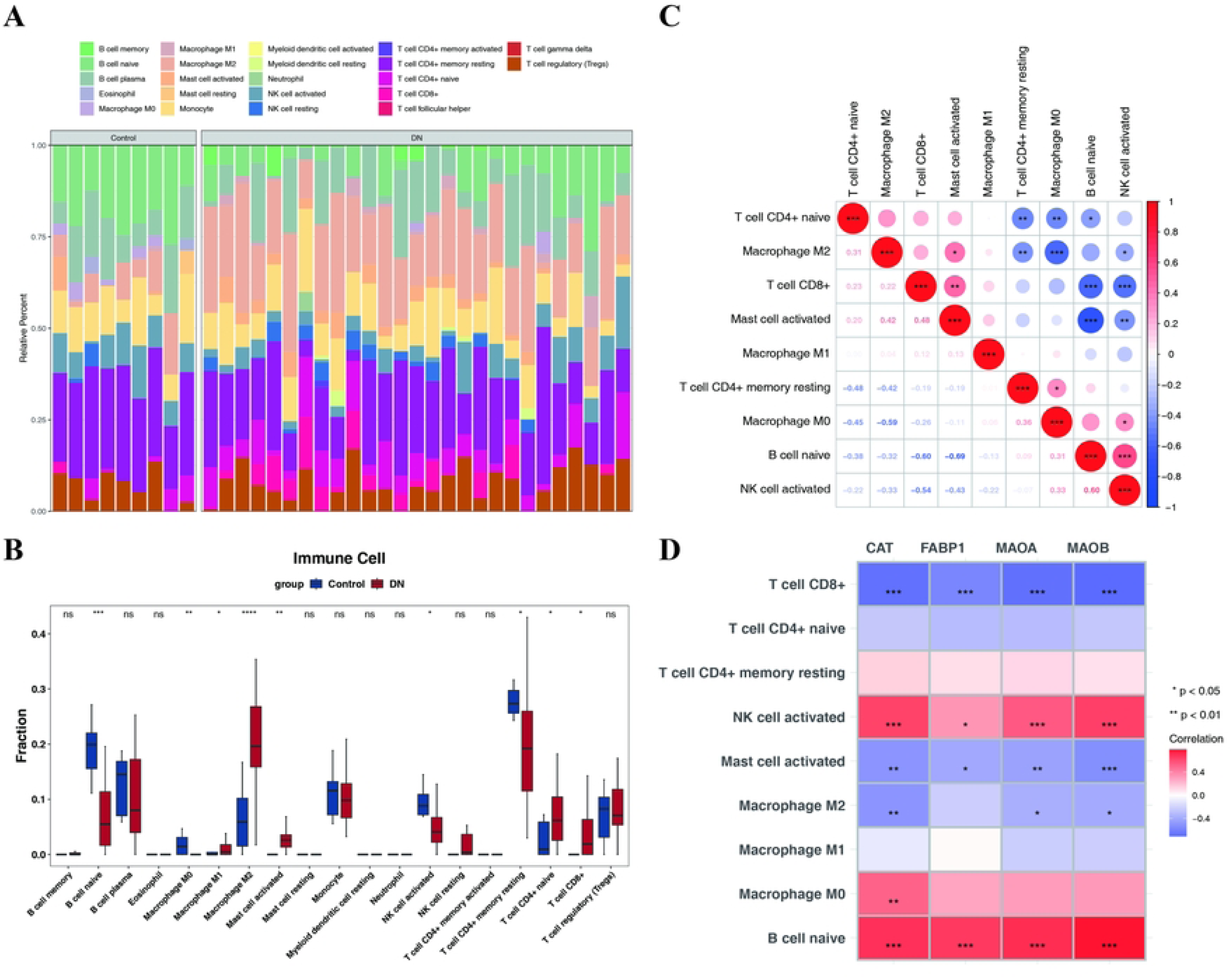
Relationship between biomarkers and immune cells. A Heatmap of differential immune cells in the DN and control groups. B Differential expression of immune cells in the DN and control groups. C Correlation of differential immune cells. D Correlation of biomarkers with differential immune cells.

### Key m6A modified proteins interacted with biomarkers

The m^6^A RNA methylation modification profoundly affects RNA synthesis and metabolism and is implicated in the pathogenesis of various diseases [30]. The locations of the m^6^A modification sites in the biomarkers and their high-confidence positions in the secondary structures are illustrated in Fig. 7A-H. Further analysis revealed that key m^6^A-modified proteins interacting with *CAT* included AQR and RBM22, while *FABP1* interacted with both SF3A3 and AQR. *MAOA* was found to interact with IGF2BP3 and IGF2BP2, and *MAOB* with TIA1 (Table 3).

**Fig. 7.**
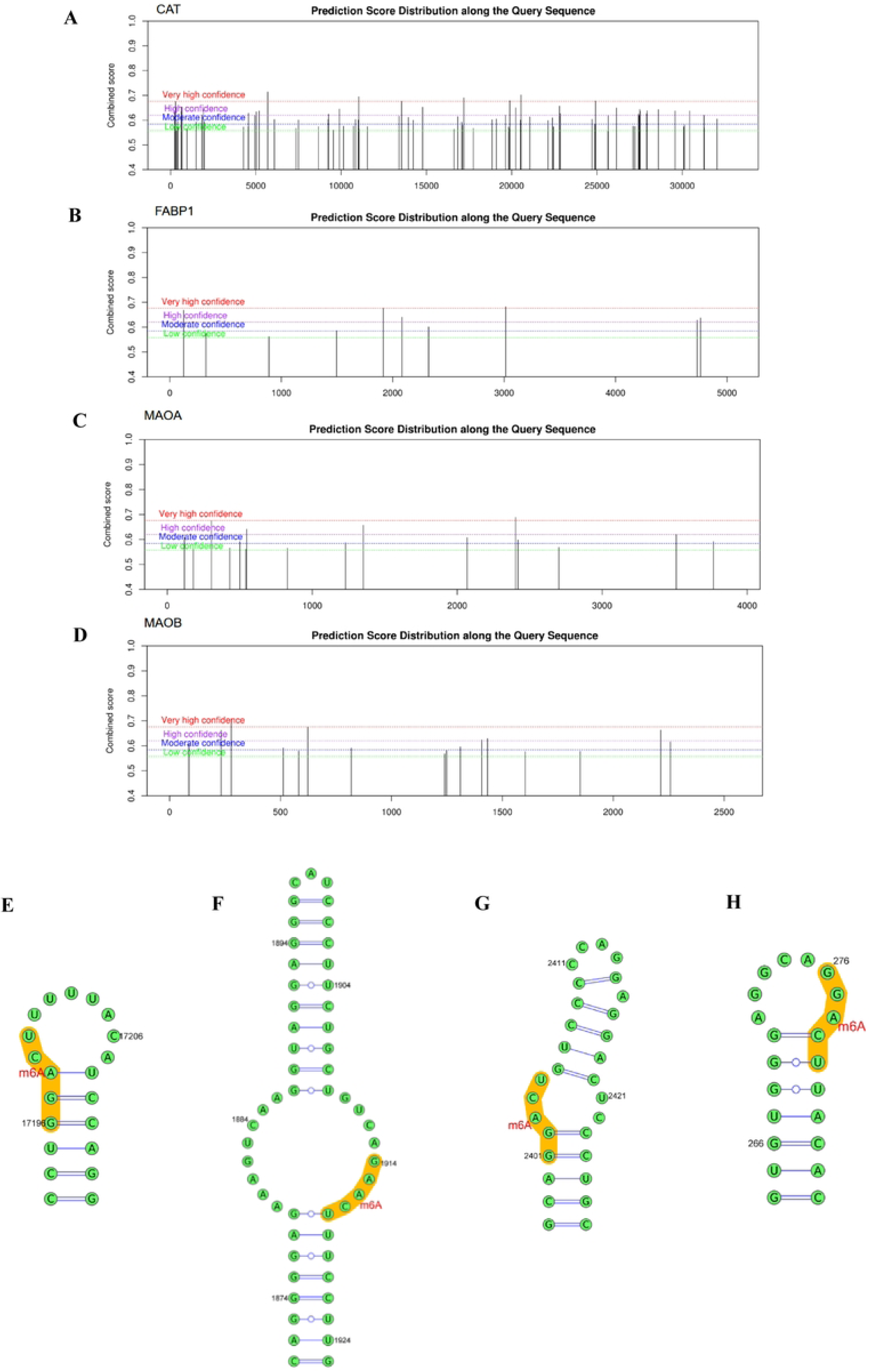
A-D The m6A modification site of the four biomarkers (CAT, FABP1, MAOA, MAOB). E-H The position of the each biomarker in the RNA secondary structure. The yellow parts represent the sequence fragments of the m6A-modified biomarkers.

**Table 3.**
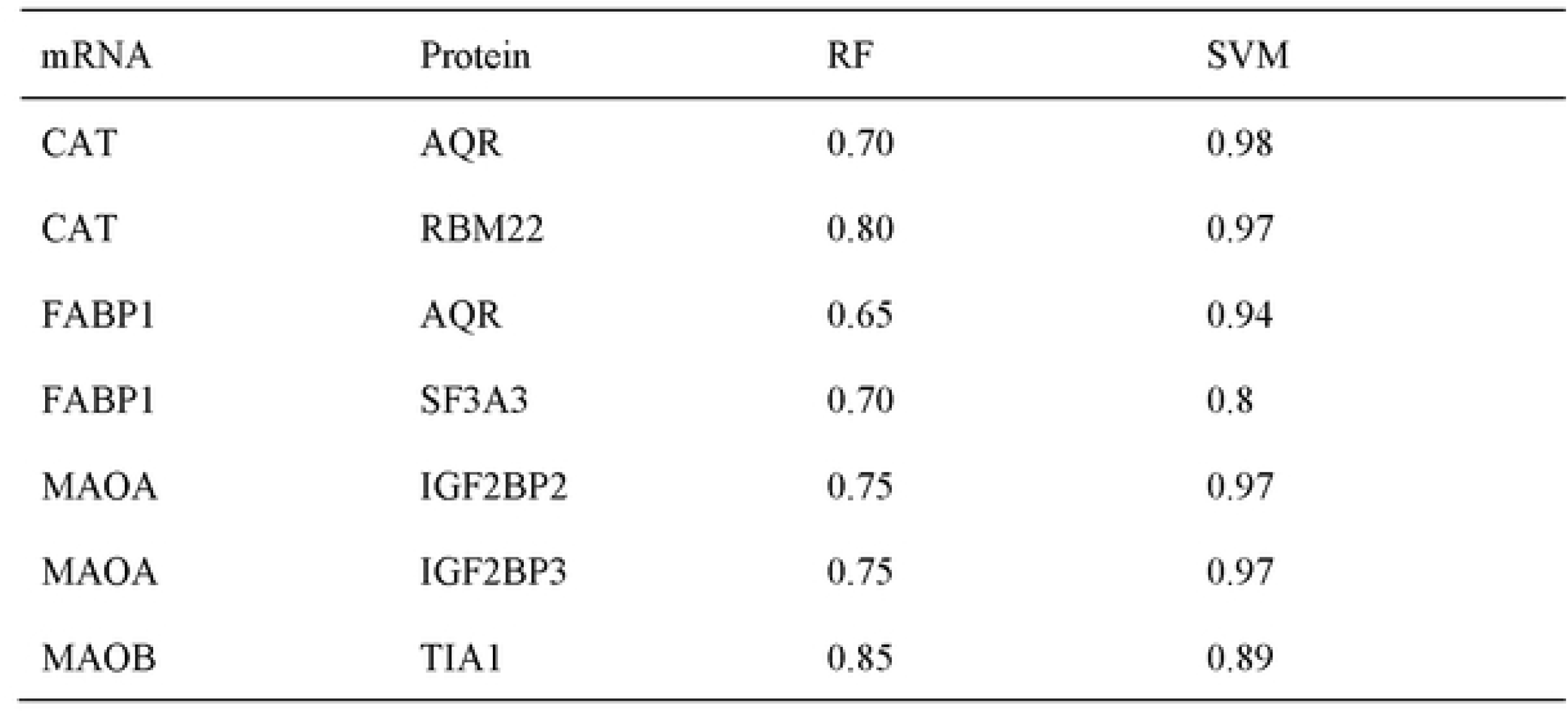
Interactions between key m6A modified proteins that interact with CAT.

| mRNA | Protein | RF | SVM |
| --- | --- | --- | --- |
| CAT | AQR | 0.70 | 0.98 |
| CAT | RBM22 | 0.80 | 0.97 |
| FABP1 | AQR | 0.65 | 0.94 |
| FABP1 | SF3A3 | 0.70 | 0.8 |
| MAOA | IGF2BP2 | 0.75 | 0.97 |
| MAOA | IGF2BP3 | 0.75 | 0.97 |
| MAOB | TIA1 | 0.85 | 0.89 |

### Taxifolin, beta-sitosterol, and eriodictyol were core active ingredient in LLF treating DN

In miRNet, *CAT* was predicted to interact with 24 miRNAs, while *FABP1* was associated with five miRNAs. Additionally, *MAOB* and *MAOA* were linked to 29 and 26 miRNAs, respectively. Among these, 23 lncRNAs were identified in both the TarBase and Starbase databases. A lncRNA-miRNA-mRNA regulatory network was then constructed, incorporating four biomarkers, 74 miRNAs, and 23 lncRNAs (Fig. 8A). Core active ingredients targeting the biomarkers included luteolin, beta-sitosterol, eriodictyol, kaempferol, quercetin, and taxifolin (Fig. 8B). Moreover, an active ingredient-biomarker-pathway network was established based on the active ingredients, biomarkers, and the top five pathways identified in GSEA (Fig. 8C). For instance, taxifolin targeted *CAT* in the peroxisome pathway. The binding energies between *CAT* and taxifolin (-8.8 kcal/mol), *FABP1* and beta-sitosterol (-8.1 kcal/mol), and *MAOB* and eriodictyol (-9.8 kcal/mol) were all below -5 kcal/mol, suggesting strong affinities between these biomarkers and their respective active ingredients [29]. Taxifolin, beta-sitosterol, and eriodictyol were identified as core active ingredients in LLF for treating DN (Fig. 8D-F).

**Fig. 8.**
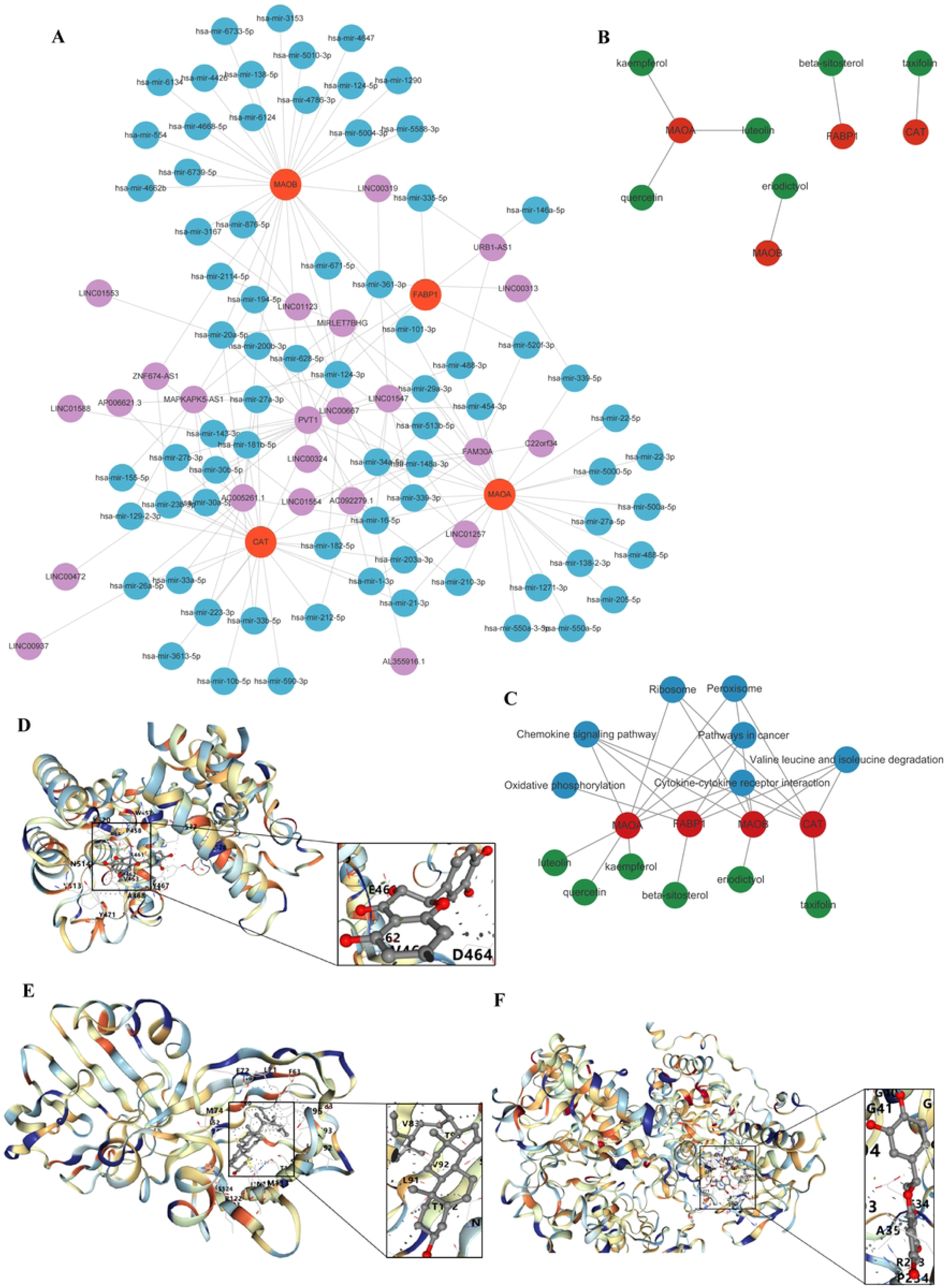
Core active ingredients of LLF. A lncRNA-miRNA-mRNA regulatory network. B Core active ingredients targeting biomarkers. C Active ingredient-biomarker-pathway network. D-F Molecular docking of CAT with taxifolin, FABP1 with beta-sitosterol, and MAOB with eriodictyol.

### Validate biomarkers in the DN mouse model

#### Pharmacodynamic evaluation of LLF in treating DN mice

During the administration period, the levels of blood glucose and urinary microalbumin in mice were dynamically monitored every two weeks (Fig. 9A-B). Compared with the control group, blood glucose and urinary microalbumin in DN model group was significantly increased (*P* < 0.01); compared with the DN model group, blood glucose of the mice in the treatment group were significantly decreased after 4 weeks of administration (*P* < 0.01) and urinary microalbumin of the mice in the treatment group were significantly decreased after 8 weeks of administration (*P* < 0.05). The results imply that LLF could be beneficial in treating DN.

**Fig. 9.**
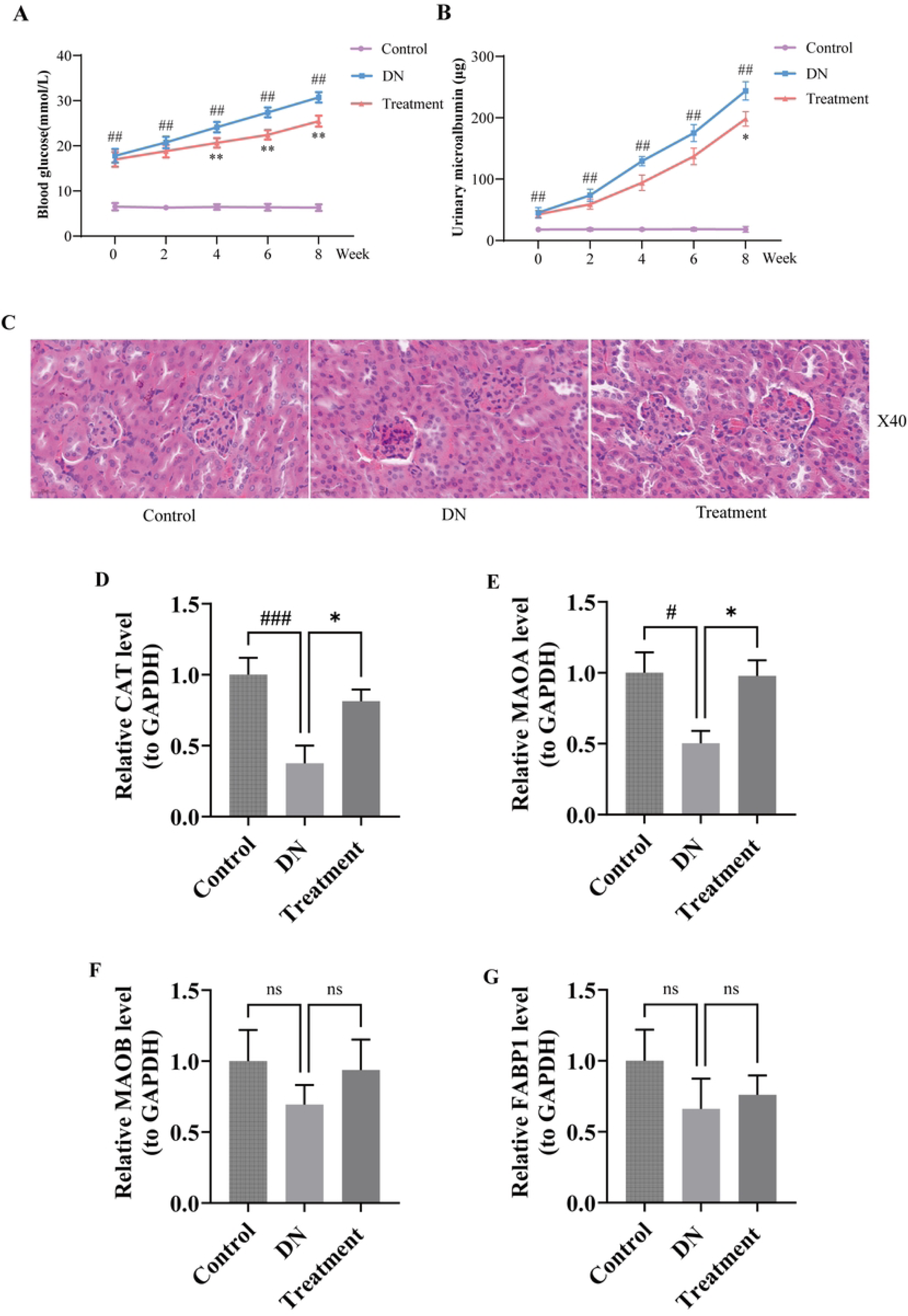
LLF reduces blood glucose levels, ameliorates pathological damage to their kidney tissues, and upregulates the expression of CAT and MAOA genes in db/db mice. A The level of blood glucose in serum of db/db mouse. B The level of urinary microalbumin in urine of db/db mouse. C Histological analysis of liver tissues by H& E staining (Magnification ×40). D-F The mRNA expression levels of CAT, MAOA, MAOB and FABP1 in renal tissue. vs control group, ^#^*P* < 0.05, ^##^*P* < 0.01, ^###^*P* < 0.001; vs DN group, \**P* < 0.05, \*\**P* < 0.001.

#### Pathological evaluation of LLF in treating DN mice

Following HE staining, the control group exhibited clear glomerular structures in kidney tissue. In contrast, the DN model group showed glomerular nuclear pyknosis and hyperchromasia, along with inflammatory cell infiltration around the glomeruli, compared to the normal group. Treatment with LLF ameliorated the pathological damage in the kidney tissue of db/db mice (Fig. 9C).

#### RT-PCR analysis the expression of biomarkers in DN mice

Following the successful establishment of a DN mouse model and the observation of symptom significant improvement with LLF treatment, RT-qPCR was further used to analyze the changes of biomarkers. Compared to the control group, the DN group exhibited significantly reduced expression of CAT and MAOA (*P* < 0.05 or *P* < 0.001). Conversely, the treatment group showed significantly increased CAT and MAOA expression compared to the DN group (*P* < 0.05). However, no statistically significant differences were observed in the expression of MAOB and FABP1 among the groups (Fig. 9D-G).

## Discussion

LLF is a widely used traditional Chinese medicine in clinical practice, primarily for nourishing the liver and kidneys [32]. It is commonly employed to treat internal heat wasting-and-thirst and is frequently prescribed for diabetes and its complications [14]. The pathogenesis of DN is influenced by various factors, including metabolic disturbances, oxidative stress, inflammatory signaling, fibrosis, and hemodynamic alterations[33]. Currently, there are no specific drugs or treatments for DN, with management relying mainly on hypoglycemic, lipid-lowering, and antihypertensive therapies. Angiotensin-converting enzyme inhibitors (ACEi) are commonly used, and in end-stage DN, renal replacement therapies such as dialysis are the only options [34–35]. However, these treatments can only slow renal damage progression in a small proportion of patients[36]. Traditional Chinese medicine, specifically LLF, has been shown to correct glucose and lipid metabolism disorders in DN rats, reducing blood glucose and lipid levels while alleviating oxidative stress in the renal cortex, thus effectively preventing DN [37]. Given that the kidney has the highest mitochondrial content and oxygen consumption after the heart, abnormalities in mitochondrial dynamics play a critical role in the pathogenesis of kidney disease [38]. This study reveals a novel mechanism by which LLF may influence mitochondrial function through specific biomarkers (*CAT*, *FABP1*, *MAOB*, and *MAOA*) in the treatment of DN.

Previous studies have indicated that catalase (*CAT*), fatty acid binding protein 1 (*FABP1*), monoamine oxidase B (*MAOB*), and monoamine oxidase A (*MAOA*) are implicated in DN to varying extents [39–40]. *CAT* is involved in the antioxidant defense system, protecting the kidney from oxidative stress-induced damage [41]. *FABP1* is involved in lipid metabolism, and its expression is closely linked to the deterioration of renal function in patients with DN. Dysregulation of *FABP1* can damage mitochondrial structure, increase the production of reactive oxygen species and malondialdehyde, and exacerbate kidney damage[42]. In other DN-related conditions, *MAOB* and *MAOA*, as enzymes involved in neurotransmitter metabolism, have been associated with the progression of DN, contributing to the imbalance in tissue redox state[43]. This study further confirms the pivotal role of these four biomarkers in DN, with their expression levels showing a decreasing trend in the DN group. It is hypothesized that modulating the expression of these biomarkers may help reduce inflammation and oxidative stress in DN.

Based on GSEA enrichment analysis, four biomarkers—*CAT*, *FABP1*, *MAOB*, and *MAOA*—were enriched in multiple pathways, including the chemokine signaling pathway, cytokine-cytokine receptor interaction, and peroxidase pathways. Chemokines are key components of the immune response, promoting inflammation [44–45]. The peroxidase (POD) pathway is linked to oxidative stress[46]. Baicalin has been reported to alleviate DN by reducing oxidative stress and inflammation, with its mechanism potentially involving the activation of the NrF2-mediated antioxidant signaling pathway and inhibition of the MAPK-mediated inflammatory pathway[48]. Moreover, dysregulation of *FABP1* in lipid metabolism may contribute to glomerular sclerosis and interstitial fibrosis in DN[9]. These findings suggest that these biomarkers play a critical role in the inflammation and oxidative stress processes of DN. Targeting these biomarkers to modulate the pathways they influence could mitigate the inflammation and oxidative stress associated with DN, thereby alleviating its progression.

In this study, bioinformatics analysis and animal experiments revealed the role of immune cell infiltration in the development of DN and the treatment of LLF. In DN, the inflammatory response is driven by a variety of cytokines and chemokines released by macrophages, lymphocytes and renal cells [49]. Bioinformatics analysis showed that the infiltration level of immune subsets such as CD8 + T cells in DN kidney tissue changed significantly, and the increase of CD8+ T cells was significantly negatively correlated with the expression of four mitochondrial-related biomarkers ( *CAT, FABP1, MAOB, MAOA* ). This calculation prediction is consistent with the pathological observation results of animal experiments : HE staining sections of the kidneys of mice in the DN model group showed obvious inflammatory cell infiltration around the glomeruli ; after LLF intervention, the infiltration of renal inflammatory cells in the treatment group was significantly reduced, and the pathological damage was improved. This suggests that increased inflammatory cell infiltration is a key feature of DN renal injury, and LLF may play a protective role by regulating immune infiltration. This finding is consistent with previous studies : the infiltration of CD8 + T cells is associated with the development of DN, and inhibition of its response can alleviate the disease[50], which also aggravates renal injury in adriamycin nephropathy [51]. In addition, a variety of immune cells such as B cells, M1 / M2 macrophages, NK cells, etc.are changed in DN pathology [52]. The biomarker CAT may affect the function of immune cells in DN [53], and MAOA may also affect the immune microenvironment by regulating macrophage polarization. These results indicate that the renal protective effect of LLF is closely related to its regulation of abnormal immune infiltration including CD8+ T cells, thereby reducing inflammatory damage.

As a natural flavonoid, Taxifolin (TA) has been shown to significantly reduce blood glucose, uric acid, creatinine, and serum insulin levels in diabetic rats, while also mitigating pathological kidney changes in these animals [54]. β-sitosterol may improve DN indirectly by regulating lipid balance and exerting anti-inflammatory effects. The β-sitosterol components in Huangqi Gegen decoction (HGD) participate in DN-related pathways, targeting molecules such as Vascular Endothelial Growth Factor A (VEGFA) and Interleukin-6 (IL-6). These effects include anti-inflammatory, anti-apoptotic, anti-oxidative, and autophagic actions, which reduce renal fibrosis and renal cortical damage and improve renal function, ultimately delaying DN progression [59]. Ophiopodiol, another natural flavonoid, has been shown to protect against ischemic stroke (IS) by balancing oxidative stress and inflammation[60]. While there are limited studies on Ophiopogon diol in the context of DN, given the disease’s association with inflammation and oxidative stress, it is hypothesized that it may alleviate DN through similar mechanisms. Drug predictions in this study also suggest that Taxifolin, β-sitosterol, and eriodictyol have potential therapeutic effects in DN.

In this study, four key biomarkers related to mitochondrial dysfunction, *CAT, FABP1, MAOB and MAOA,* were successfully identified by integrating transcriptome data and LLF active component information, and their roles in LFF treatment of DN were revealed. These markers showed good discrimination ability in diagnostic performance verification ( AUC > 0.70 ), suggesting their potential as diagnostic indicators for DN. The nomogram model based on the above markers can be used for DN risk prediction. Further functional enrichment, immune infiltration, regulatory network and molecular docking analysis preliminarily clarified the mechanism of these markers in DN. Animal experiments showed that LLF intervention could significantly improve hyperglycemia and renal pathological damage in db / db mice, and up-regulate the expression of CAT and MAOA in renal tissue. The results confirmed that the expression of the four markers decreased in DN samples, and the difference in the expression of CAT and MAOA was statistically significant ( P < 0.05 or P < 0.001 ). These findings provide new insights into the potential mechanism of LLF in the treatment of DN, and also lay a theoretical foundation for the development of DN therapeutic targets for mitochondrial dysfunction. However, this study is mainly based on bioinformatics and network pharmacology analysis. Some results ( such as m6A modification analysis, regulatory network and molecular docking ) are still limited to database prediction, and have not been further verified by methylation sequencing and cell function experiments. In the future, more in-depth experimental studies and large-scale clinical trials are still needed to clarify the specific mechanism of action and clinical transformation value of these biomarkers.

## Acnkowledgements

We would like to express our sincere gratitude to all individuals and organizations who supported and assisted us throughout this research.

## Author contributions

Qinqing Li: Investigation, Formal analysis, Writing - review & editing, Data curation. Yang Jie: Investigation, Data curation. Yanli Xin: Investigation. Lixia Wen: Investigation. Kaiwen Li: Software, Data curation. Ruqiao Luan: Investigation, Conceptualization. Xuelan Zhang: Investigation, Supervision. All authors have reviewed the results and approve the final version of the manuscript.

## Ethics statement

Ethical approval for this experiment was obtained from Shanxi University of Chinese Medicine (2022DW167).

